# Genome of the parasitoid wasp *Dinocampus coccinellae* reveals extensive duplications, accelerated evolution, and independent origins of thelytokous parthenogeny and solitary behavior

**DOI:** 10.1101/2021.06.30.450623

**Authors:** Arun Sethuraman, Alicia Tovar, Walker Welch, Ryan Dettmers, Camila Arce, Timothy Skaggs, Alexander Rothenberg, Roxane Saisho, Bryce Summerhays, Ryan Cartmill, Christy Grenier, Yumary Vasquez, Hannah Vansant, John Obrycki

## Abstract

*Dinocampus coccinellae* (Hymenoptera: Braconidae) is a generalist parasitoid wasp that parasitizes >50 species of predatory lady beetles (Coleoptera: Coccinellidae), with thelytokous parthenogeny as its primary mode of reproduction. Here we present the first high quality genome of *D. coccinellae* using a combination of short and long read sequencing technologies, followed by assembly and scaffolding of chromosomal segments using Chicago+ HiC technologies. We also present a first-pass ab initio genome annotation, and resolve timings of divergence and evolution of (1) solitary behavior vs eusociality, (2) arrhenotokous vs thelytokous parthenogenesis, and (3) rates of gene loss and gain among Hymenopteran lineages. Our study finds (1) at least two independent origins of eusociality and solitary behavior among Hymenoptera, (2) two independent origins of thelytokous parthenogenesis from ancestral arrhenotoky, and (3) accelerated rates of gene duplications, loss, and gain along the lineages leading to *D. coccinellae*. Our work both affirms the ancient divergence of Braconid wasps from ancestral Hymenopterans and accelerated rates of evolution in response to adaptations to novel hosts, including polyDNA viral co-evolution.

## Introduction

Hymenopterans are an iconic group among the diverse and species rich insect orders and encompass expansive species radiations across sawflies, ants, bees and wasps, dating back to the Carboniferous and Triassic periods 329–239 million years ago [mya] (Branstetter et al., 2018; Gauld, 1988; Malm and Nyman, 2015; Peters et al., 2017). Current consensus across contemporary studies into resolving the phylogeny of Hymenoptera establish the divergence of sawfly and woodwasp lineages (‘Symphyta') from all other hymenopterans (‘Apocrita’) near the basal branches in this order’s evolutionary history, while Apocrita further radiated into a diverse range of parasitoids (‘Parasitica’ clade) and stinging insects (‘Aculeata’ clade) which comprises stinging wasps, bees and ants (Branstetter et al., 2018; Gauld, 1988; Malm and Nyman, 2015; Vilhelmsen and Turrisi, 2011). Amongst the many evolutionary novelties to arise among Hymenopterans, are differential modes of reproduction (e.g., sexual, thelytokous and arrhenotokous parthenogenesis), ecto- and endo-parasitism, and eusocial behavior. Recent work suggests that the most recent common ancestor [MRCA] of Hymenoptera was phytophagous and originally consumed living plant tissues (Peter et al., 2017). Therefore, the transition from phytophagy to parasitism has been hypothesized to radiate from a “single endophylytic parasitoid” wasp predecessor between 289-211 mya in the Permian or Triassic period (Peter et al., 2017). Specifically, among Braconid wasps, the Euphorinae subfamily predominantly exploits the adult stage of their hosts (koinobiosis), which is not the most common mode of host resource exploitation relative to the majority of parasitoid wasps (Stigenberg et al., 2015). It has also been posited that ancestral members of the Euphorine clade may have shifted host-resource exploitation from ovipositing within juvenile hosts to adult hosts which were in the same location, allowing for further adaptive radiations to their host (Quicke et al., 1990).

*Dinocampus coccinellae* (Hymenoptera: Braconidae-Euphorinae) is a parthenogenetic, generalist parasitoid wasp with a cosmopolitan distribution, observed to parasitize over fifty species of lady beetles (Coleoptera: Coccinellidae) (Ceryngier et al., 2017) across the world. Characteristic of other Braconid wasps, parasitoid larvae feed on their insect hosts throughout development until they eclose as an an adult female ready to oviposit unfertilized eggs into a host (Shaw *et al., 1991*). However, unlike other endoparasitoids in the Euphorinae subfamily that are largely koinobionts with characteristically narrow host ranges (Shaw *et al.*, 1991), *D. coccinellae* are generalist endoparasitoids that parasitize incognito (endophytic) hosts (Ceryngier *et al*., 2017). *D.coccinellae* is observed to predominantly reproduce parthenogenetically (thelytokous), with the rare occurrence of observed males in the population (Wright, 1979). Little is known about their evolutionary history, or host-shifting tactics, with some recent work indicating considerable phenotypic plasticity in size-morphology of emergent daughter wasps (Vansant et al., 2019) covarying with the size of their hosts. Mutations and chromosomal segregation should therefore account for all genetic variation in each new generation of mostly clonal *D. coccinellae* (Slobodchikoff and Daly, 1971), with no recombination.

*D. coccinellae* are also solitary wasps, with limited interactions between other conspecific individuals, unlike eusocial wasps. Eusociality and solitary behavior have been long proposed to have independently evolved among Hymenopteran lineages (Hines et al., 2007; Kuhn et al., 2019). Eusociality was previously interpreted to have singular origins in vespid wasps, deriving eusocial behavior from a singular common ancestor in Hymenoptera (Morale et al., 2004). However, through multi-gene phylogeny analyses, it has been observed that eusociality may have evolved twice in vespid wasps (Hines et al., 2007). Nonetheless, the origins of solitary behavior in Hymenoptera are yet to be delineated, primarily owing to the absence of genomic data among solitary wasps. Another interesting aspect of *D. coccinellae*’s biology involves individual wasps harboring an RNA virus (D*inocampus coccinellae* Paralysis Virus, DcPV) that replicate in the cerebral ganglia cells of their coccinellid hosts, thereby manipulating their behavior (Dheilly et al., 2015). This endosymbiotic parasitic relationship between *D. coccinellae* and DcPV thus suggests the independent co-evolution of genes involved in antiviral response, and host behavioral manipulation, with accelerated gene-family evolution among genes involved in host-parasite conflicts.

As a first attempt to address many of these questions and to decipher the evolutionary history of *D. coccinellae*, here we sequence the first high-resolution genome of the species, followed by a first-pass annotation and phylogenomic analysis of *D. coccinellae* in the context of other Hymenopterans sequenced as part of the i5K project. Our analyses provide the foundation for future research in understanding the genomics of host-shifts, behavioral manipulation, and parthenogenesis in a unique parasitoid wasp species.

## Methods

### Samples, Wasp Rearing

Parthenogenetic lines of female *D. coccinellae* that were collected from the field in Summer 2018 were raised on a laboratory population of *Coccinella septempunctata* from Kansas, USA (JJO pers. comm.), and *Hippodamia convergens* obtained from Green Thumb Nursery, San Marcos, CA, USA. Stocks of *C. septempunctata* and *H. convergens* were maintained in separate insect tents (fed on pea aphids *ad libitum* – *Acyrthosiphon pisum*, raised on fava bean plants – *Vicia faba*) in a greenhouse at CSUSM, San Marcos, California. Parthenogenetic lines (>20 individual wasps) were collected from exposing female *D. coccinellae* to multiple host beetles over their lifetime, and thereon flash-frozen using liquid nitrogen, and maintained at –80C until further processing. Genomic DNA was then extracted using the Qiagen DNeasy Kit, following the manufacturer’s protocols. DNA quality was then assessed using a 1% agarose gel and quantified using a Qubit 2.0 Fluorometer with broad range standards.

### Chicago(R) library preparation and sequencing (Dovetail Genomics)

The protocols of Putnam et al., 2016 and Erez Lieberman-Aiden et al., 2009 were then used to produce a Chicago(R) library and a Dovetail HiC library respectively. Briefly ~500ng of quality-assessed, high-molecular weight genomic DNA was subject to chromatin reconstitution *in vitro*, then fixed with formaldehyde. Fixed chromatin was then digested with the DpnII (NEB), 5’ overhangs filled in with biotinylated nucleotides, and then free blunt ends were ligated. After ligation, crosslinks were reversed and the DNA purified from protein. Purified DNA was treated to remove biotin that was not internal to ligated fragments. The DNA was then sheared to ~350 bp mean fragment size and sequencing libraries were generated using NEBNext Ultra enzymes and Illumina-compatible adapters. Biotin-containing fragments were isolated using streptavidin beads before PCR enrichment of each library. The Chicago and HiC libraries were then sequenced on an Illumina HiSeq X at Dovetail Genomics.

### PacBio Library and Sequencing

The manufacturer recommended protocol was used to generate a PacBio SMRTbell library (~20kb) for PacBio Sequel using SMRTbell Express Template Prep Kit 2.0 (PacBio, Menlo Park, CA, USA). The library was bound to polymerase using the Sequel II Binding Kit 2.0 (PacBio) and loaded onto PacBio Sequel II. Sequencing was then performed on PacBio Sequel II 8M SMRT cells at Dovetail Genomics, generating 179 Gb of raw data.

### De Novo Genome Assembly

The long-read assembler, Wtdbg2 (Ruan and Li, 2020) was used to assemble the genome (--genome_size 0.2g --read_type sq --min_read_len 5000). Blobtools v1.1.1 (Laetsch and Blaxter) was used to identify potential contamination in the assembly based on NCBI BLAST (v2.9) hits of the assembly against the NT database. A fraction of the scaffolds was identified as contaminants and were removed from the assembly. The filtered assembly was then used as an input to purge_dups v1.1.2 (Guan et al. 2020), and potential haplotypic duplications were removed from the assembly.

### Scaffolding with Chicago and HiC HiRise

The input de novo assembly after filtering for contaminations and duplicate haplotypes, Chicago library reads, and Dovetail HiC library reads were used as input to Dovetail’s HiRise, a software pipeline designed specifically for using proximity ligation data to scaffold genome assemblies (Putnam et al., 2016). An iterative analysis was then conducted, comprising the following steps: (1) Chicago library sequences were aligned to the draft input assembly using a modified SNAP read mapper (http://snap.cs.berkeley.edu), (2) the separations of Chicago read pairs mapped within draft scaffolds were analyzed by HiRise to produce a likelihood model for genomic distance between read pairs, and the model was used to identify and break putative misjoins, to score prospective joins, and to make joins above a threshold, and (3) after aligning and scaffolding the Chicago library reads, Dovetail HiC library sequences were aligned and scaffolded following the same method. Quality of these final scaffolded assemblies were assessed using N50, N90 and other genome continuity statistics, prior to additional bioinformatic analyses.

### Ab initio gene prediction, Repeat Masking

AUGUSTUS v.3.3.3 (Keller et al., 2011) was used to predict protein coding genes and coding sequences on the final HiRise genome assembly, using the *Nasonia vitripennis* genome annotation as a training set (Rago et al., 2016). Repeat masking was also performed on the HiRise assembly using RepeatMasker v.4.0.9 (Smit et al., 2019). The *Drosophila melanogaster* family in the Dfam v.3.3 library was used as a reference repeat library, and all output annotations were obtained as GFF3 formatted files.

### Ortholog Identification, Core gene completeness

All amino acid sequences predicted by AUGUSTUS were then uploaded to OrthoDB v.10.1 and orthologous amino acid sequences were identified using five Hymenopteran genomes - *Microplitis demolitor*, genome GCF_000572035.2, *Nasonia vitripennis*, genome GCF_000002325.3, *Neodiprion lecontei*, genome GCF_001263575.1, *Orussus abietinus*, genome GCF_000612105.2, and *Trichogramma pretiosum*, genome GCF_000599845.2. Completeness of the HiRise assembly was assessed using BUSCO v.5.0 (Seppey et al., 2019), against core genes from all Eukaryotes (eukaryota_odb10.2019-11-20 - 255 BUSCO markers), Insects (insecta_odb10.2019-11-20 - 1367 BUSCO markers), and Hymenoptera (hymenoptera_odb10.2019-11-20 - 5991 BUSCO markers).

### Multiple Sequence Alignment, Species Tree Reconstruction

A BLAST database was then constructed using the AUGUSTUS predicted gene-set, and the complete list of identified orthologs for *D. coccinellae* was then “intersected” with the list of single copy amino acid sequences from the i5K project (Thomas et al., 2020) by using BLASTP and obtaining the scaffold coordinates across the *D. coccinellae* genome. Separate FASTA files (for each orthologous single copy gene) were then constructed with all the i5K Hymenopteran genomes and our *D. coccinellae* genome, and multiple sequence alignments constructed using pasta v.1.8.6 (Mirarab et al., 2015). RAxML v.8.2.12 (Stamakis 2014) was then used to construct gene trees using the PROTGAMMAJTTF amino acid substitution model, *sensu* Thomas et al., 2020. ASTRAL v. 5.7.7 (Zhang et al., 2018) was then utilized to infer an unrooted species tree.

### Time Calibration, Ancestral State Reconstruction

The species tree obtained from ASTRAL was then time-calibrated using the fossil-times derived from Thomas et al., 2020 (common ancestor of *Athalia rosae* and all other hymenopterans – 226.4-411 mya, common ancestor of Formicidae (ants), and Anthophila (bees) - 89.9-93.9 mya, common ancestor of Apis (honeybees) and Bombus (bumblebees), Melipona (stingless bees) - 23-28.4 mya). 95 random orthologous amino acid locus alignments were concatenated from across the 26 species analyzed (with *Zootermopsis nevadenisis* as outgroup) and analyzed using the approximate likelihood method implemented in mcmctree (Yang 2007). Briefly, the estimation of divergence times and branch lengths is conducted in two steps: (1) branch lengths are estimated using a maximum likelihood method, and (2) divergence times are then estimated using an MCMC method. The root-age was set to be < 1000 mya, and likelihood estimation was performed using the JC69 model, followed by a long MCMC run (2e7 iterations discarded as burn-in, followed by 1e7 iterations, sampling every 10 iterations, generating a total of 1e6 samples). Convergence of the MCMC was then assessed using Tracer 1.7.1 (Rambaut et al., 2018) by observing the traces of all divergence time parameter estimates, and ESS values. The time-calibrated rooted tree obtained from mcmctree was then used for ancestral state reconstruction using the phytools package in R (Revell 2012). Specifically, we used the (a) discrete state reconstruction, and (b) empirical Bayes reconstruction using 1000 simulated trees for two relevant Hymenopteran traits - (a) mode of reproduction – thelytoky (unfertilized eggs developing into females), arrhenotoky (unfertilized eggs developing into males), and sexual reproduction, (b) sociality – solitary, eusociality, and facultative sociality.

### Gene Family Evolution

All protein coding gene sequences from Hymenoptera from the study of Thomas et al., 2020 were obtained from www.arthrofam.org and together with the *ab initio* protein predictions from our AUGUSTUS run, were parsed through the OrthoFinder pipeline (Emms and Kelley 2019) to perform comparative genomic analyses of (a) gene duplications, (b) identifying single copy orthologs, and (c) delineating orthogroups based on reciprocal DendroBLAST/DIAMOND searches and estimating gene-trees. The gene family counts identified by OrthoFinder and a rooted, binary, and ultrametric species tree (based on the species tree inferred above) were then used in iterative runs of the likelihood-based method, CAFE5 (Mendes et al., 2020) to estimate gene turnover rates (λ) and annotation error rates (ε), sensu the methods of Thomas et al., 2020.

Additionally, a parsimony method (DupliPHY v.1.0 – Ames and Lovell 2015) was used to obtain accurate ancestral gene counts. Significant rapid evolution (gene gain or loss) was then assessed by regressing gene counts at internal nodes (ancestral) versus external (extant) nodes, with statistical significance assessed at > 2 standard deviations of the variance within the gene family.

## Results

### Genome Assembly Quality and Completeness

The final HiRise assembly from Dovetail Genomics suggests an approximate genome size of 182 Mbp in 720 scaffolds, with a total of 183 gaps, an N50 of 8.6 Mbp, and N90 score of 536 Kbp. The largest scaffold was 19 Mbp, with 99.72% of scaffolds > 1 Kbp in length, and contained an average of 10.05 missing base-calls (N’s) per 100 kbp. Our assembly of *D. coccinellae* is thus by far the most complete, and contiguous of all publicly available parasitoid wasp genomes in the i5K project (http://i5k.github.io/arthropod_genomes_at_ncbi). Analyses of BUSCO completeness using the eukaryota_odb10 database obtained 94.9% completeness (242 out of 255 groups searched), with >89% completeness upon comparison with the insecta_odb10 and hymenoptera_odb10 databases. Identification of repeats and transposable elements with RepeatMasker with the Dfam v.3.3 *Drosophila melanogaster* database identified a total of 690 retroelements (~191 kbp), 89 DNA transposons (~11 kbp) and other simple repeat and satellite regions (see Table 3). A comprehensive annotation of repeats thus identified 8.84% (~16 Mbp) of the genome to be comprised of repeats.

**Table 1:** Genome contiguity statistics from across Braconid wasp genomes that are publicly available, in comparison with the high-quality *D. coccinellae* genome of this study. Of note are the comparable genome sizes (126.92-384.37 Mbp), and % GC content (30.6-39.4%). Our *D. coccinellae* assembly however presents a several-fold improvement in contig and scaffold N50.

| Species | Source | Genome Size (Mbp) | Contig N50 (bp) | Scaffold N50 (bp) | % GC |
| --- | --- | --- | --- | --- | --- |
| *Dinocampus coccinellae* | this study | 182.09 | 8604445 | 8604445 | 38.77 |
| *Cotesia vestalis* | i5k | 186.1 | 46055 | 46055 | 30.6 |
| *Diachasma alloeum* | i5k | 384.37 | 45480 | 657001 | 38.9 |
| *Fopius arisanus* | i5k | 153.63 | 51867 | 978588 | 39.4 |
| *Macrocentrus cingulum* | i5k | 127.92 | 65089 | 65089 | 35.6 |
| *Microplitis demolitor* | i5k | 241.19 | 14116 | 1139389 | 33.1 |

**Table 2:**
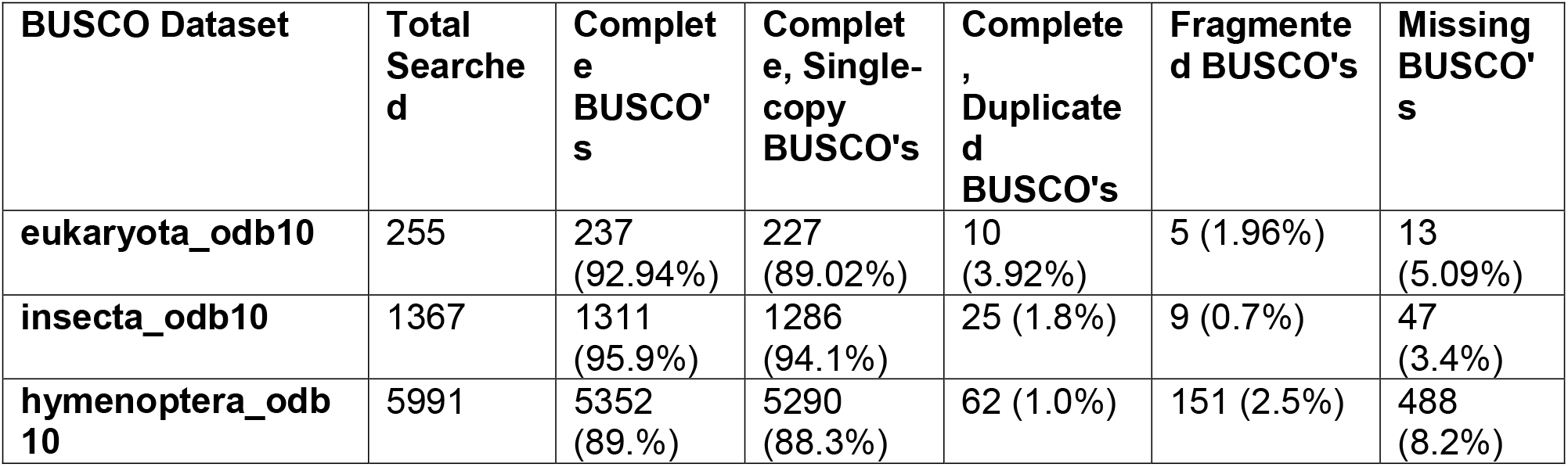
BUSCO genome completeness measures of the *D. coccinellae* HiRise assembly, when compared to three separate databases: (1) eukaryota, (2) insecta, and (3) hymenoptera. BUSCO completeness assess the quality of a genome assembly by mapping core genes and gene families that are highly conserved across taxa.

**Table 3:**
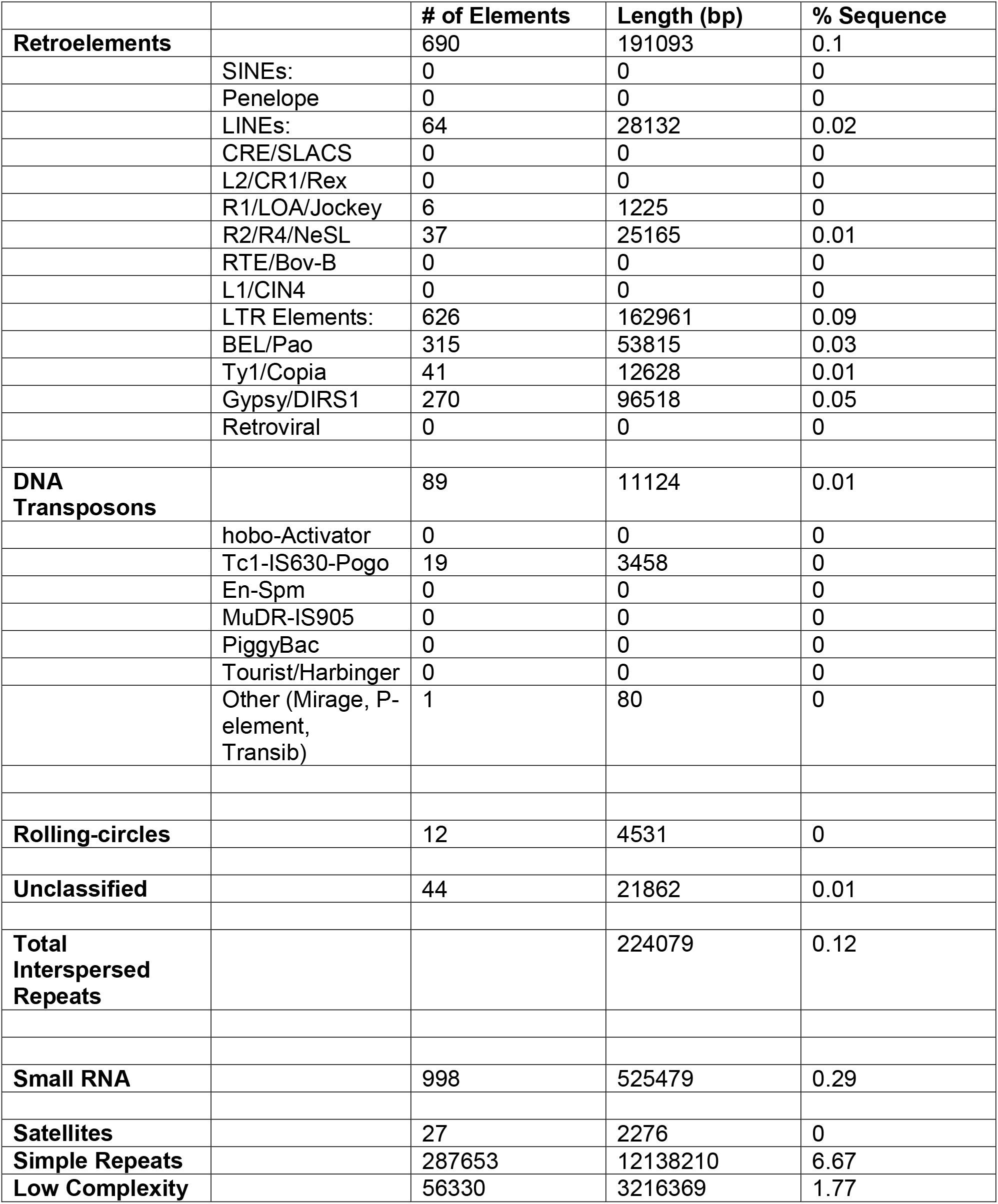
Assessment of retroelements, transposable elements, and other repeats across the *D. coccinellae* HiRise assembly using RepeatMasker v.4.0.9 (Smit et al., 2019), with further annotations from RepeatMasker obtained as a GFF3 track that can be visualized on the genome annotation using JBrowse.

### Genome Annotation and Orthology

Ab initio gene prediction using AUGUSTUS v.3.3.3 (Keller et al., 2011) identified a total of 68,797 protein coding sequences in the HiRise assembly. All gene annotations were then added as a separate track to create a genome browser (JBrowse) instance, which can be accessed at www.github.com/arunsethuraman/dcoccinellae.

Orthology prediction using OrthoDB v.10.0 against five other parasitoid wasp genomes obtained over 8000 orthogroups (longest – EGF-like calcium binding domain 5at7399, and Immunoglobulins 0at7399). Annotations for these orthologs were obtained and corresponding GO terms associated were catalogued.

### Phylogeny, Time Calibration and Ancestral State Reconstruction

Phylogeny reconstruction of the species tree using ASTRAL from 2045 gene trees placed *D. coccinellae* as sister to other parasitoid wasps (*Trichogramma pretiosum*, *Copidosoma floridanum,* and *Nasonia vitripensis*). The remainder of the tree replicated the same species tree topology obtained from ASTRAL and RAxML analyses from Thomas et al., 2020, which resolves wasps as sister to the common ancestor of all ants and bees.

Fossil-based time calibration of the ASTRAL species tree obtained above with MCMCtree, and utilizing 200 randomly sampled genes (out of 2045) determined the split of the outgroup (*Zootermopsis nevadensis)* and all hymenopterans at 561 mya (95% HPD interval of 256.93 mya – 954.48 mya), the split of *D. coccinellae* from other wasps (*T. pretiosum, C. floridanum*, *N. vitripennis)* at 135.88 mya (95% HPD interval of 108.87 mya – 166.69 mya), and the split of Apidae (bees) and Formicidae (ants) at 91.94 mya (95% HPD interval of 89.92 mya – 93.91 mya). Ancestral state reconstruction of mode of reproduction using phytools with the MCMC method of Huelsenbeck et al., 2003 (stochastic character mapping) revealed independent evolution of thelytoky along the branches leading to *D. coccinellae* and *T. pretiosum,* with their common ancestors determined to have been arrhenotokous. Similarly, sexual reproduction was determined to have evolved independently in the common ancestor of *A. cephalotes* and *A. echinatior.* A similar estimation of ancestral state reconstruction for sociality estimated the independent convergent evolution of eusociality within the Hymenoptera along at least three lineages: (1) within Apidae, in the common ancestor of *B. terrestris*, *B. impatiens*, *M. quadrifasciata*, *A. mellifera*, and *A. florea*, (2) in the common ancestor of all Formicidae, and (3) along the branch leading to *C. floridanum*. Interestingly, the common ancestor of all bees, ants, and wasps was determined to have exhibited predominantly solitary behavior, with facultative sociality evolving independently along the *E. mexicana* lineage.

### Gene Family Evolution

Discovery of orthologous sequences using OrthoFinder with 25 Hymenoptera genomes (24 from i5k, and *D. coccinellae*) determined 96.1% of all genes analyzed assigned to 19,210 unique orthogroups, 3116 of which contained all species, and 1241 contained single-copy orthogroups. Interestingly, *D. coccinellae* was determined to have a total of 40,012 gene duplication events, several folds larger than other hymenopterans (nearly 10-fold higher than *N. vitripennis,* with 4180 gene duplication events). CAFE5 estimated gene turnover rate (λ) of 0.145 (with a maximum possible λ of 0.41). DupliPHY analyses to determine accelerated rates of gene loss or gain across all single-copy ortholog families analyzed determined several significant gene loss events along the *D. coccinellae* lineage (see Supplement …). The most significant gene loss events included genes in families of olfactory/odorant receptors, membrane proteins, and uncharacterized helix-turn-helix motifs across Hymenoptera. Gene gain events spanned families of transposases (e.g. harbinger), endonucleases involved in stress response in the Bombus lineages, and membrane transport proteins (e.g. carboxyl transferases).

## Discussion

Here, we present the first high quality genome of the thelytokous parasitoid wasp, *Dinocampus coccinellae*, a species known for its unique solitary life history cycle, RNA viral-mutualism, and plasticity in parasitism across host coccinellid species. Our analyses indicate (a) ancient divergence of *D. coccinellae* from ancestral parasitoid wasps (~136 MYA), (b) extensive gene duplications (~10x more than *Nasonia vitripennis*), (c) multiple independent evolutionary shifts to solitary behavior among Hymenoptera, (d) at least two independent shifts from ancestral arrhenotoky to thelytoky, and (e) accelerated evolution among several gene families along the *D. coccinellae* lineage.

The phylogeny of Braconid wasps are yet to be delineated using whole genomes, with most of the current work utilizing mitochondrial genes, and morphological information (Chen and Achterberg, 2019) to inform origins and divergence. Here we utilize a phylogenomic approach to delineate an ancient divergence of Braconidae (here represented by *D. coccinellae)* from the common ancestor of other parasitoid wasps in the Jurassic-Cretaceous period (~136 MYA). Braconid wasps are speciose, with a variety of endo- and ecto-parasitic life history strategies adapted to adult hosts among Coleoptera, Hemiptera, and Lepidoptera, with several independently evolved novel polydnavirus mutualisms (Herniou et al., 2013). Our work affirms the timeline proposed by Herniou et al., 2013 using polydnavirus genomes, and lays the foundation for understanding models of viral-parasitoid wasp-host coevolution and diversification. These timelines are also in lines with previous work that establishes the timing of evolution of the three major modes of sex determination and reproduction across Hymenoptera: sexual reproduction, arrhenotokous parthenogenesis, and thelytokous parthenogenesis. Arrhenotokous parthenogenesis (arrhenotoky) has been determined to be the ancestral mode of sex determination and reproduction dating back to as far as 300 mya, and presently remains the most prominent mode throughout the Hymenoptera order; arrhenotoky describes the process of sex determination in which diploid females develop from fertilized eggs and unfertilized eggs give rise to haploid males (Beukeboom et al., 2007; Heimpel and Jetske, 2008; Slobodhcikoff and Daly, 1971). Thelytokous parthenogenesis (thelytoky) on the other hand is a convergently derived mode of sex determination and reproduction in which diploid female wasps are born from unfertilized egg clones (Beukeboom et al., 2007; Heimpel and Jetske, 2008; Slobodhcikoff and Daly, 1971; Kuhn et al., 2019).

Similarly, among Ichneumonoid parasitoid wasps, it is established that this apocritan superfamily consists of two main subfamilies: the Braconidae and Ichneumonidae sister clades (Belshaw et al., 2002; Quicke et al., 2020) with differential parasitism modes. Across these two sister branches, the different host parasitism strategies that parasitoid wasps employ center around how they exploit various developmental stages of their host. The first host exploitation strategy, idiobiosis, describes parasitoids that oviposit into immobilized hosts with paused development during the larval parasitoid’s growth, such as host eggs or cocooned juveniles; contrastingly, koinobiosis, describes parasitoids which oviposit into adult or larval hosts that continue to eat and develop further throughout larval parasitoid growth (Belshaw et al., 1998; Harvey et al., 2016; Jervis et al., 2011). Generally, it has been determined that the host resource exploitation strategy employed most often by the Ichneumonidae subfamily tends to favor idiobiont ectoparasitoids, while the Braconidae subfamily often exploit their hosts through a koinobiont endoparasitoid strategy (Gauld, 1988; Quicke et al., 1990). Between the Braconidae and Ichneumonidae sister subfamilies, ancestral members of both branches similarly externally exploit their hosts juvenile/immature stages (idiobiont ectoparasitoid), then going on to radiate across a range of different hosts over time (Gauld, 1988). Our study therefore also affirms this timeline of adaptive evolution to host-parasitism.

Contemporary analysis into the origins of thelytokous parthenogeny (unfertilized eggs develop into daughters) in Hymenoptera point to this reproductive strategy having convergently evolved from an ancestral arrhenotokous haplo-diploid state (Kuhn et al., 2019), with the conditional utilization of sexual reproduction to “restore” genetic diversity. However, the high morphological variability of D. *coccinellae,* despite the presence of a clonal genome, may suggest that the restoration of genetic diversity is unnecessary given their immense ability to change with their environment (Vansant et al., 2019). Ancestral state reconstruction in this study points to at least two independent evolutionary events leading to thelytoky among parasitoid wasps (Fig.), which also interestingly coincides with the evolution of solitary behavior in *D. coccinellae* and *T. pretiosum* (Fig.). Solitary behavior and thelytoky can be seen as complimentary behaviors as the absence of mates encourages asexual reproduction.

The evolution of eusociality is just one of many major transitions on Earth (Woodard et al. 2011). The transition to eusocial from solitary has occurred many times, mostly in insects, and only in a small number of lineages (Woodard et al. 2011). The evolution of eusociality is quite interesting since it requires a balance between cooperation and conflict with a preferential shift towards cooperation since this would be the only favorable outcome for fitness (Woodard et al. 2011). The selective pressures for individual success, such as in the case of the Braconid wasp *Dinocampus coccinellae,* requires that the amount of energy put into the offspring outweighs the costs of forgoing reproduction to care for the offspring of others, as in the case of the A*pis mellifera* where the wellbeing of the hive is one of the top priorities (Woodard et al. 2011).

Evidence for the influence of environmental factors suggests that relatedness and kinship may also play an important role in the development of eusociality (Hughes et al., 2008). However, there is also evidence for sociality determination through environmental factors (Soucy & Danforth, 2002). Recent research of Hymenopteran chemoreceptors and their vast differentiation and specialization among different species have shown to play a major role in the emergence and development of eusocial behavior (Ferguson et al., 2021). Chemoreceptor genes and their frequencies are highly variable among Hymenopteran species, generally occurring in large expansions of eusocial species, but are also known to have lineage-specific patterns of losing or gaining genes due to tandem repeat events that result in unique clusters of chemoreceptor genes (Ferguson et al., 2021). Our genome-wide analyses of diversification and gene gain/loss events finds consistent gene loss and gain among chemoreceptor and viral-coevolution genes along the *D. coccinellae* lineage. Further work utilizing gene-expression analyses are required to establish the functional significance of these genes among Hymenoptera.

## Acknowledgments

This work was funded by the National Institute of Food and Agriculture, U.S. Department of Agriculture, Hatch Program under accession number 1008480 and funds from the University of Kentucky Bobby C. Pass Research Professorship to JJO, NSF ABI Development #1664918 to AS, USDA NIFA #2017-06423 to PI George Vourlitis and co-PI AS, NSF REU #1852189 to PI Betsy Read and co-PI AS, CSUSM GPSM #86969 to AS, and a CSUPERB COVID-19 Research Recovery Microgrant to AS. This research includes calculations carried out on HPC resources supported in part by the National Science Foundation through major research instrumentation grant number 1625061 and by the US Army Research Laboratory under contract number W911NF- 16-2-0189. Special thanks to Ryan Ames (Exeter) and Ben Fulton (Indiana) for their help with the DupliPHY and CAFE5 analyses respectively. AS would also like to acknowledge Marc Tollis, Jeet Sukumaran, Kent Holsinger, Liam Revell, Milton Tran, and other “phylogenetics tweeps” for their invaluable recommendations on species-tree inference and ancestral state reconstruction. AS also graciously acknowledges Cianna Bedford-Petersen for developing the R color palette based on the Super-Women of the US Presidential Inauguration of 2021 (https://github.com/ciannabp/inauguration).

---

**Fig. 1.**
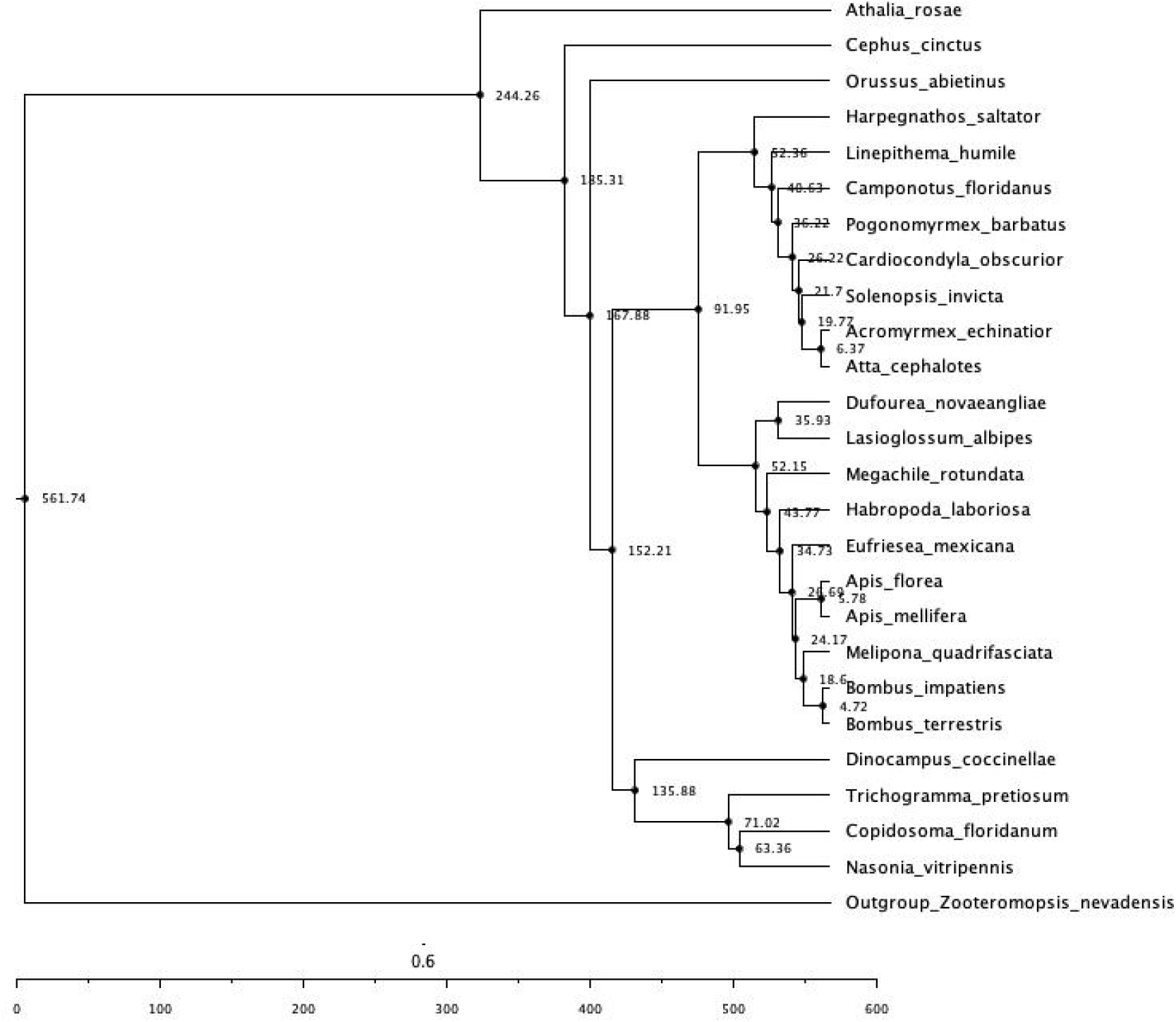
Fossil-time calibrated species tree using 200 random genes (from 2045 total genes) across all publicly available hymenopteran genomes in the i5k Project, with *Dinocampus coccinellae* placed as being sister to other Braconid wasps. All nodes are in units of million years ago (mya), and branch lengths are scaled by time, as indicated by the scale below.

**Fig. 2:**
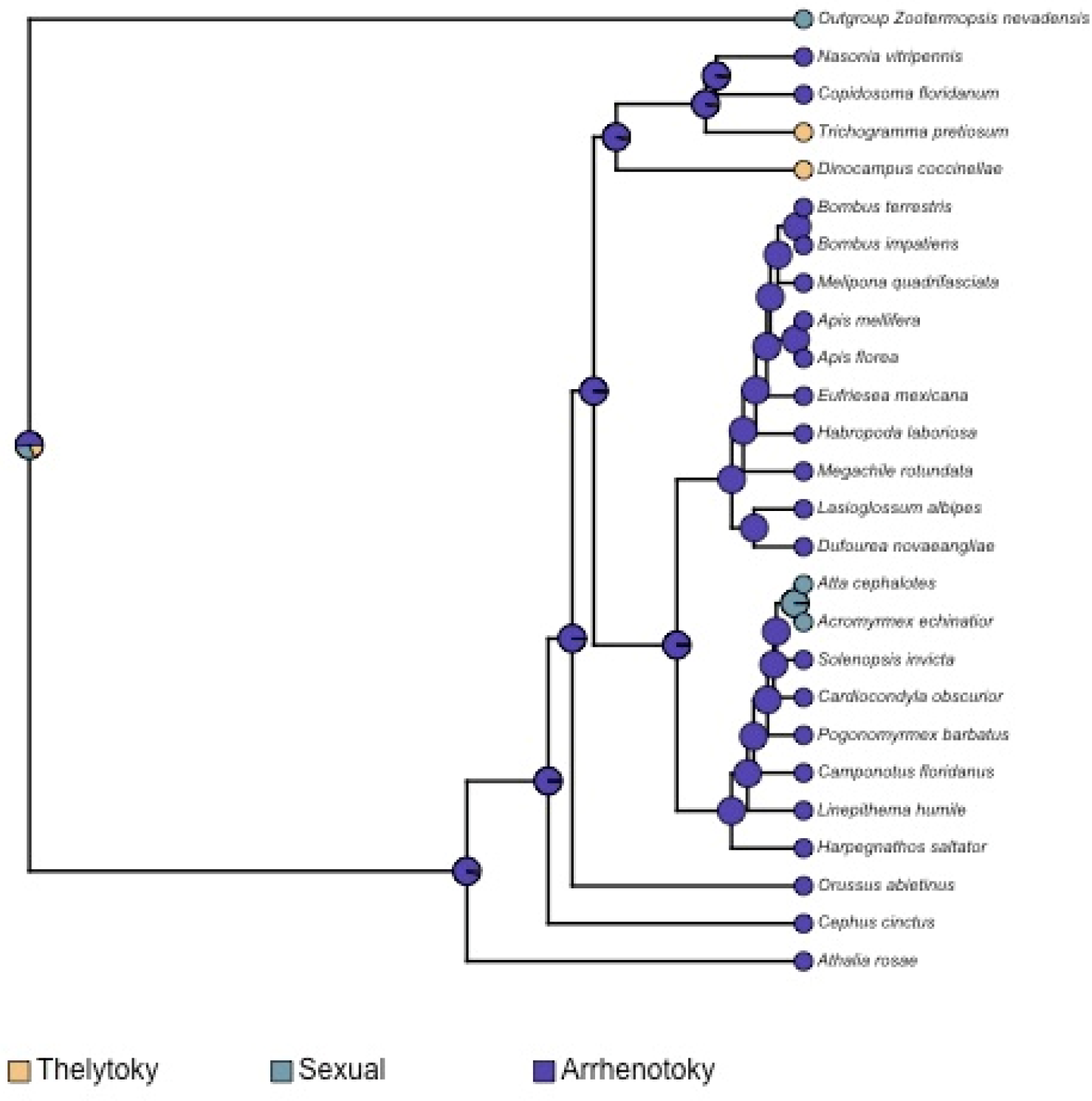
Ancestral state reconstruction using the stochastic character mapping method of Huelsenbeck et al., 2003, as implemented in phytools (Revell 2012), mapping the evolution of mode of parthenogenesis across all hymenopterans, in comparison with the outgroup, *Zootermopsis nevadensis.* The pies at internal and external nodes represent the posterior probability distribution of one of three possible states: (1) Thelytoky, (2) Arrhenotoky, and (3) Sexual reproduction.

**Fig. 3:**
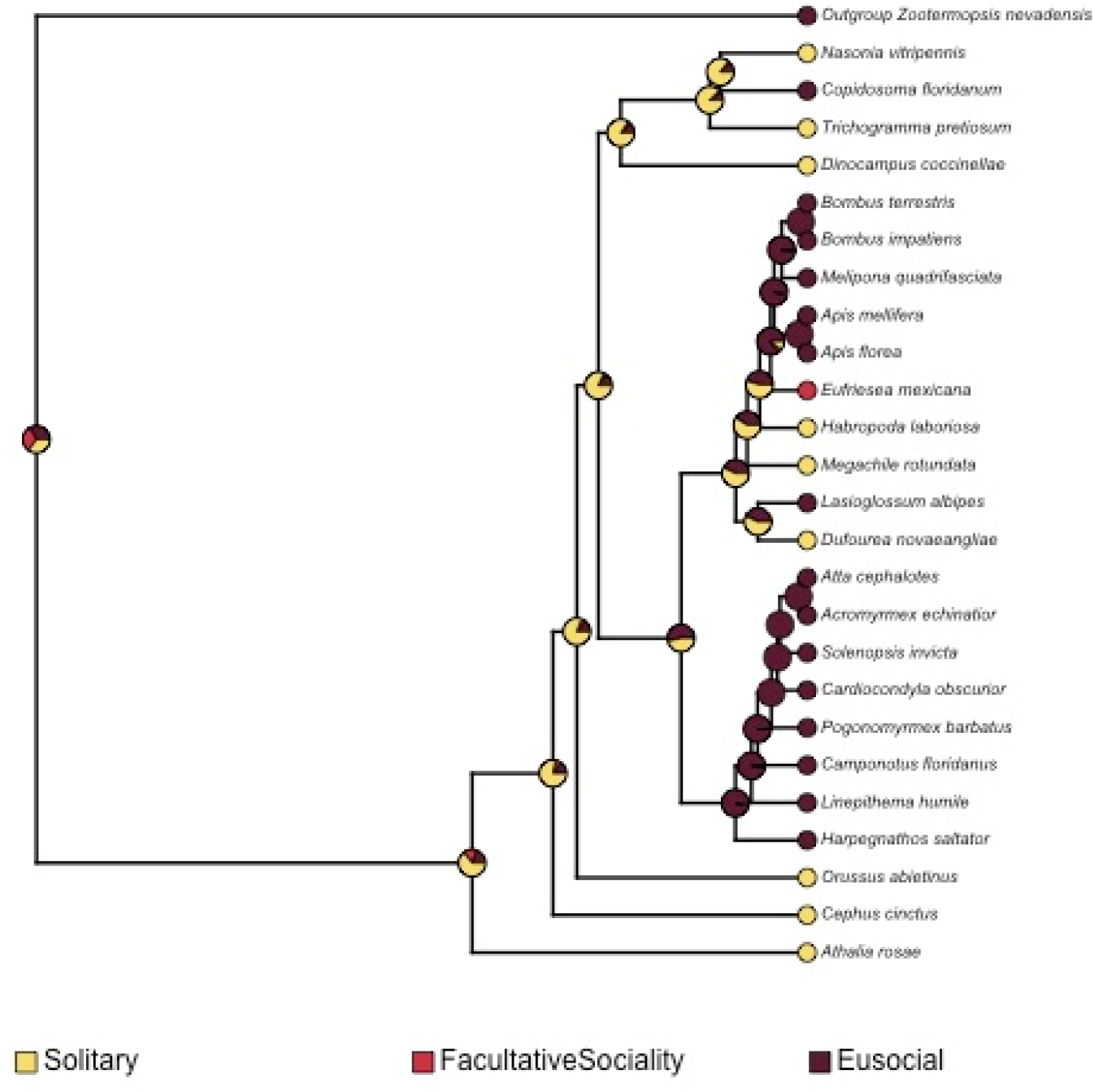
Ancestral state reconstruction using the stochastic character mapping method of Huelsenbeck et al., 2003, as implemented in phytools (Revell 2012), mapping the evolution of sociality across all hymenopterans, in comparison with the outgroup, *Zootermopsis nevadensis.* The pies at internal and external nodes represent the posterior probability distribution of one of three possible states: (1) Solitary, (2) Eusociality, and (3) Facultative sociality.

## References

Belshaw, R., & Quicke, D. L. (2002). Robustness of ancestral state estimates: evolution of life history strategy in ichneumonoid parasitoids. Systematic Biology, 51(3), 450–477.

Beukeboom, L. W., Kamping, A., & van de Zande, L. (2007, June). Sex determination in the haplodiploid wasp Nasonia vitripennis (Hymenoptera: Chalcidoidea): a critical consideration of models and evidence. In Seminars in cell & developmental biology (Vol. 18, No. 3, pp. 371–378). Academic Press.

Branstetter, M. G., Childers, A. K., Cox-Foster, D., Hopper, K. R., Kapheim, K. M., Toth, A. L., & Worley, K. C. (2018). Genomes of the Hymenoptera. Current opinion in insect science, 25, 65–75.

Ceryngier, P., Nedvěd, O., Grez, A. A., Riddick, E. W., Roy, H. E., San Martin, G., … & Haelewaters, D. (2018). Predators and parasitoids of the harlequin ladybird, Harmonia axyridis, in its native range and invaded areas. Biological Invasions, 20(4), 1009–1031.

Chen, Xue-xin, and Cornelis van Achterberg. “Systematics, phylogeny, and evolution of braconid wasps: 30 years of progress.” Annual Review of Entomology 64 (2019): 335–358.

Dheilly, N. M., Maure, F., Ravallec, M., Galinier, R., Doyon, J., Duval, D., … & Mitta, G. (2015). Who is the puppet master? Replication of a parasitic wasp-associated virus correlates with host behaviour manipulation. Proceedings of the Royal Society B: Biological Sciences, 282(1803), 20142773.

Erez Lieberman-Aiden, Nynke L. van Berkum, Louise Williams, Maxim Imakaev, Tobias Ragoczy, Agnes Telling, Ido Amit, Bryan R. Lajoie, Peter J. Sabo, Michael O. Dorschner, Richard Sandstrom, Bradley Bernstein, M. A. Bender, Mark Groudine, Andreas Gnirke, John Stamatoyannopoulos, Leonid A. Mirny, Eric S. Lander, Job Dekker. Comprehensive Mapping of Long-Range Interactions Reveals Folding Principles of the Human Genome. Science. 2009; 326, 289.

Ferguson, S. T., Ray, A., & Zwiebel, L. J. (2021). Olfactory genomics of eusociality within the Hymenoptera. In Insect Pheromone Biochemistry and Molecular Biology (pp. 507–546). Academic Press.

Gauld, I. D. (1988). Evolutionary patterns of host utilization by ichneumonoid parasitoids (Hymenoptera: Ichneumonidae and Braconidae). Biological Journal of the Linnean Society, 35(4), 351–377.

Guan, D., McCarthy, S. A., Wood, J., Howe, K., Wang, Y., & Durbin, R. (2020). Identifying and removing haplotypic duplication in primary genome assemblies. Bioinformatics, 36(9), 2896–2898.

Harvey, J. A., & Malcicka, M. (2016). Nutritional integration between insect hosts and koinobiont parasitoids in an evolutionary framework. Entomologia experimentalis et applicata, 159(2), 181–188.

Heimpel, George E., and Jetske G. De Boer. “Sex determination in the Hymenoptera.” Annu. Rev. Entomol. 53 (2008): 209–230.

Herniou, E. A., Huguet, E., Thézé, J., Bézier, A., Periquet, G., & Drezen, J. M. (2013). When parasitic wasps hijacked viruses: genomic and functional evolution of polydnaviruses. Philosophical Transactions of the Royal Society B: Biological Sciences, 368(1626), 20130051.

Hines, H. M., Hunt, J. H., O’Connor, T. K., Gillespie, J. J., & Cameron, S. A. (2007). Multigene phylogeny reveals eusociality evolved twice in vespid wasps. Proceedings of the National Academy of Sciences, 104(9), 3295–3299.

Huelsenbeck, J. P., Nielsen, R., & Bollback, J. P. (2003). Stochastic mapping of morphological characters. Systematic biology, 52(2), 131–158.

Hughes, W. O., Oldroyd, B. P., Beekman, M., & Ratnieks, F. L. (2008). Ancestral monogamy shows kin selection is key to the evolution of eusociality. Science, 320(5880), 1213–1216.

Jervis, M., & Ferns, P. (2011). Towards a general perspective on life-history evolution and diversification in parasitoid wasps. Biological Journal of the Linnean Society, 104(2), 443–461.

Kuhn, A., Darras, H., Paknia, O., & Aron, S. (2020). Repeated evolution of queen parthenogenesis and social hybridogenesis in Cataglyphis desert ants. Molecular ecology, 29(3), 549–564.

Malm, T., & Nyman, T. (2015). Phylogeny of the symphytan grade of Hymenoptera: new pieces into the old jigsaw (fly) puzzle. Cladistics, 31(1), 1–17.

Mendes, F. K., Vanderpool, D., Fulton, B., & Hahn, M. W. (2020). CAFE 5 models variation in evolutionary rates among gene families. Bioinformatics, 36(22–23), 5516–5518.

Mirarab S, Nguyen N, Guo S, Wang L-S, Kim J, Warnow T. PASTA: Ultra-Large Multiple Sequence Alignment for Nucleotide and Amino-Acid Sequences. J Comput Biol. 2015;22(5):377–386. doi:10.1089/cmb.2014.0156.

Peters, R. S., Krogmann, L., Mayer, C., Donath, A., Gunkel, S., Meusemann, K., … & Niehuis, O. (2017). Evolutionary history of the Hymenoptera. Current Biology, 27(7), 1013–1018.

Putnam NH, O’Connell BL, Stites JC, Rice BJ, Blanchette M, Calef R, Troll CJ, Fields A, Hartley PD, Sugnet CW, Haussler D, Rokhsar DS, Green RE. Genome Research. 2016 Mar;26(3):342–50.

Quicke, D. L., Austin, A. D., Fagan‐Jeffries, E. P., Hebert, P. D., & Butcher, B. A. (2020). Molecular phylogeny places the enigmatic subfamily Masoninae within the Ichneumonidae, not the Braconidae. Zoologica Scripta, 49(1), 64–71.

Quicke, D. L., & van Achterberg, C. (1990). Phylogeny of the subfamilies of the family Braconidae (Hymenoptera: Ichneumonoidea). Nationaal Natuurhistorisch Museum.

Rago, A., Gilbert, D. G., Choi, J. H., Sackton, T. B., Wang, X., Kelkar, Y. D., … & Colbourne, J. K. (2016). OGS2: genome re-annotation of the jewel wasp Nasonia vitripennis. BMC genomics, 17(1), 1–25.

Rambaut, A., Drummond, A. J., Xie, D., Baele, G., & Suchard, M. A. (2018). Posterior summarization in Bayesian phylogenetics using Tracer 1.7. Systematic biology, 67(5), 901.

Revell, L. J. (2012). phytools: an R package for phylogenetic comparative biology (and other things). Methods in ecology and evolution, 3(2), 217–223.

Ruan, J., & Li, H. (2020). Fast and accurate long-read assembly with wtdbg2. Nature methods, 17(2), 155–158.

Seppey, M., Manni, M., & Zdobnov, E. M. (2019). BUSCO: assessing genome assembly and annotation completeness. Methods in molecular biology (Clifton, NJ), 1962, 227–245.

Shaw, M. R., & Huddleston, T. (1991). Classification and biology of braconid wasps (Vol. 7, No. 11). Royal Entomological Society.

Slobodchikoff, C. N., & Daly, H. V. (1971). Systematic and evolutionary implications of parthenogenesis in the Hymenoptera. American Zoologist, 11(2), 273–282.

Smit, A. F. A., Hubley, R., & Green, P. (2019). 2013–2015. RepeatMasker Open-4.0.

Soucy, S. L., & Danforth, B. N. (2002). Phylogeography of the socially polymorphic sweat bee Halictus rubicundus (Hymenoptera: Halictidae). Evolution, 56(2), 330–341.

Stamatakis, A. (2014). RAxML version 8: a tool for phylogenetic analysis and post-analysis of large phylogenies. Bioinformatics, 30(9), 1312–1313.

Stigenberg, J., Boring, C. A., & Ronquist, F. (2015). Phylogeny of the parasitic wasp subfamily Euphorinae (Braconidae) and evolution of its host preferences. Systematic Entomology, 40(3), 570–591.

Thomas, G. W., Dohmen, E., Hughes, D. S., Murali, S. C., Poelchau, M., Glastad, K., … & Richards, S. (2020). Gene content evolution in the arthropods. Genome biology, 21(1), 1–14.

Vansant, H., Vasquez, Y. M., Obrycki, J. J., & Sethuraman, A. (2019). Coccinellid host morphology dictates morphological diversity of the parasitoid wasp Dinocampus coccinellae. Biological Control, 133, 110–116.

Vilhelmsen, L., & Turrisi, G. F. (2011). Per arborem ad astra: Morphological adaptations to exploiting the woody habitat in the early evolution of Hymenoptera. Arthropod Structure & Development, 40(1), 2–20.

Wright, E. J. 1979. Observations on the copulatory behaviour of Perilitus coccinellae (Hymenoptera: Braconidae). Proceedings of the Entomological Society of Ontario 109: 22.

Yang, Z. (2007). PAML 4: phylogenetic analysis by maximum likelihood. Molecular biology and evolution, 24(8), 1586–1591.

Zhang, C., Rabiee, M., Sayyari, E., & Mirarab, S. (2018). ASTRAL-III: polynomial time species tree reconstruction from partially resolved gene trees. BMC bioinformatics, 19(6), 15–30.

